# Sleeping ORANGE: A Transposase-Enhanced CRISPR Approach to Boost Endogenous Protein Tagging Efficiency

**DOI:** 10.1101/2025.10.23.684145

**Authors:** Emily-Rose Martin, Josan G. Martin, Kaiyven A. Leslie, Mark A. Russell, Asami Oguro-Ando

## Abstract

**Background:** Investigating the subcellular distribution of proteins is crucial for understanding complex cell behaviours and disease mechanisms, and fluorescence microscopy has become a key tool for visualising protein localisation. Endogenous protein tagging, where the sequence for a tag (typically a peptide or fluorescent protein) is integrated into the native genetic sequence encoding a protein of interest, enables proteins to be visualised without the need for antibodies against the target protein. ORANGE (Open Resource for the Application of Neuronal Genome Editing) is a CRISPR-Cas9-based endogenous protein tagging technique which relies on homology-independent targeted integration (HITI)-mediated gene editing. Utilising HITI as the DNA repair pathway of choice gives ORANGE the advantage of being more efficient than classical homology-directed repair (HDR)-based endogenous protein tagging techniques and additionally, means it can be used in post-mitotic cells.

**Results:** We applied the ORANGE system to tag three proteins, CYFIP1, JAKMIP1, and STAT3, and confirmed that the expressed fusion proteins demonstrate expected subcellular localisations through fluorescence microscopy. Unexpectedly, the efficiency of ORANGE editing was less than 1% in HEK293 cells, despite high transfection efficiency. To improve the editing efficiency associated with ORANGE, we combined the ORANGE method with an established Sleeping Beauty transposase/CRISPR-Cas9 fusion technique, which has been shown to enhance HITI-mediated gene editing. Using this new method, which we term Sleeping ORANGE, we successfully tagged CYFIP1 with the fluorescent protein mNeonGreen. Importantly, quantitative analysis by fluorescence microscopy and flow cytometry demonstrated an increase in editing efficiency using Sleeping ORANGE, with an approximately 12.85-fold increase in the percentage of mNeonGreen-expressing cells at 72 hours post-transfection relative to populations of cells edited with the ORANGE method.

**Conclusions:** We have incorporated the DNA-binding domain of the Sleeping Beauty transposase to create a new system that improves the gene-editing efficiency of the ORANGE technique. With further developments to optimise CRISPR gRNA design and reduce off-target effects, the Sleeping ORANGE technique may form a valuable tool for researchers to better understand subcellular localisation and dynamics.

## Introduction

Investigating the subcellular localisation and dynamic functions of proteins is important for understanding disease mechanisms and to identify novel therapeutic targets (1). This can be achieved by a wide variety of techniques and visualisation of a protein of interest typically relies on immunostaining. However, immunostaining techniques have several disadvantages, particularly due to reliance on the availability of high-quality antibodies, which limits the protein targets that can be studied (2). Moreover, although immunostaining of fixed cells and tissues can be used to investigate subcellular localisation of a specific protein at a particular point in time, this technique does not reveal insights about protein dynamics occurring in real-time. Therefore, to understand the dynamic processes happening in cells, techniques compatible with live cells are better suited. Previously, studying proteins in live cells, for example through live-cell imaging experiments, often relied upon electroporating labelled antibodies, which can be a harsh and cytotoxic process (3); or inducing the overexpression of recombinant proteins tagged with a reporter marker via plasmid vectors (1). However, the use of strong transcriptional promoters in such plasmids to force expression of the target protein, along with the lack of regulatory elements necessary for normal gene expression, often produces artefacts, such as the mis-localisation of target proteins or altered cellular morphology (2,4,5).

To overcome the limitations associated with immunostaining or overexpression of recombinant proteins, researchers can instead tag the endogenously expressed proteins. Endogenous protein tagging uses gene-editing techniques to insert a tag into the genetic sequence of a protein, thereby enabling the protein to be visualised in live or fixed cells (6). Several types of tags can be inserted into the gene, such as fluorescent proteins (7), small peptide tags (8), or luminescent tags (9), depending on the purpose of the experiment. Given that endogenous protein tagging does not involve overexpressing proteins of interest, it has the advantage that proteins are expressed within their natural regulatory context, meaning that findings from these studies are likely more relevant to normal physiology (10,11). Moreover, endogenous tagging enables proteins to be visualised or quantified even when no validated antibodies against them are available (2) and enables visualisation of dynamic events with a short lifetime (9).

There are several established methods for endogenously tagging proteins of interest, the majority of which utilise clustered regularly interspaced short palindromic repeats (CRISPR)-Cas9 gene editing technologies. Open Resource for the Application of Neuronal Genome Editing (ORANGE) is a CRISPR-Cas9 based gene editing technique, but rather than using homology-directed repair (HDR) to insert a donor DNA sequence, ORANGE utilises homology-independent targeted integration (HITI) (2). HITI is based on the non-homologous end-joining (NHEJ) pathway of DNA repair and has been demonstrated to be much more efficient than HDR mechanisms, and because it is based on NHEJ, HITI can be used for gene editing in post-mitotic cells (12). ORANGE utilises a plasmid vector encoding a CRISPR guide RNA (gRNA) targeting the gene of interest, along with *Streptococcus pyogenes*-derived Cas9 (spCas9) and a donor DNA template to facilitate HITI (2). The donor DNA template is constructed of the fluorescent or small peptide tag sequence flanked by gRNA and downstream protospacer adjacent motif (PAM) sequences (2). Cleavage by Cas9 creates blunt ends on the donor DNA template and within the target gene, thereby enabling the donor DNA to be inserted into the target gene cleavage site by NHEJ (2). Willems *et al*. used ORANGE to tag an array of neuronal proteins with green fluorescent protein (GFP), showing that ORANGE accurately tags endogenous proteins without affecting the localisation of the target proteins (2).

In this study, we use ORANGE to tag three proteins with three fluorescent tags: Cytoplasmic and FMRP-interacting Protein 1 (CYFIP1), a cytoplasmic protein that plays roles in regulating neuronal translation and Actin remodelling, with mNeonGreen; Janus Kinase and Microtubule-Interacting Protein 1 (JAKMIP1), an RNA-binding protein that stabilises microtubules, with mScarlet; and Signal Transducer and Activator of Transcription 3 (STAT3), a transcription factor activated downstream of cytokine receptor activation, with mGold. Through fluorescence microscopy, we demonstrate that the expressed fusion proteins show the expected subcellular localisations, but that the gene-editing efficiency of the endogenous tagging is very low. We then develop a novel strategy for endogenous protein tagging based on the ORANGE technique by combining it with the DNA-binding capabilities of the Sleeping Beauty transposase. As proof-of-concept, we successfully used this novel method, which we term Sleeping ORANGE, to tag CYFIP1 and found that it improved ORANGE gene-editing efficiency.

## Methods and Materials

### gRNA design

A gRNA sequences for *CYFIP1* (5’-CGCGTCCTCCAGAGTCACCT-3’), *JAKMIP1* (5’-GTCGAAGAAAGGCCGGAGCA-3’) and *STAT3* (5’-GCTGCTGTAGCTGATTCCAT-3’) were designed using the CRISPR tool available in the *Benchling* software (Benchling [Biology Software] (2024)) and purchased from *Integrated DNA Technologies* (along with their complementary sequence) with sticky ends to facilitate type IIS restriction enzyme cloning using BbsI.

### ORANGE donor tag design

The donor tags for the ORANGE constructs were designed using the *Benchling* software, using the gRNAs and fluorescent protein sequences (mNeonGreen for CYFIP1, mScarlet for JAKMIP1 and mGold for STAT3) obtained from *FPbase* (13) and *SnapGene*® (from Dotmatics; available at snapgene.com). Where necessary to prevent frameshifts in the DNA sequence and to improve the flexibility of the protein structure, linker sequences were added to the donor tag sequences based on commonly published ‘safe’ linker sequences (14). The donor tags were purchased from *Integrated DNA Technologies*, in the form of a custom gene in a vector flanked with the HindIII and XhoI restriction sites to facilitate cloning (referred to as pORANGE tag construct (pOR TagC) plasmids).

### Sleeping ORANGE donor tag design

The donor tag for the *CYFIP1* ORANGE construct was modified using the *Benchling* software to design the Sleeping ORANGE version of the *CYFIP1* donor tag. The canonical Sleeping Beauty-binding DNA sequence (referred to as the N57 binding sequence) based on (15) was adapted and inserted into the donor tag between the first gRNA sequence and the sequence encoding mNeonGreen. Adaptations to the N57 binding sequence included shortening of the inverted repeat (IR) sequences to reduce the size of the N57 binding sequence and modification of premature stop codons within the IR sequences to prevent nonsense-mediated decay of the modified mRNA sequence. Additionally, the sequence encoding a Puromycin resistance gene was inserted downstream of the adapted N57 binding sequence to enable selection of successfully transfected and edited cells. A P2A self-cleaving peptide sequence was added directly before the mNeonGreen start codon to ensure that the additional peptide produced from the N57 binding sequence did not affect the structure of the translated CYFIP1-mNeonGreen protein. Finally, the entire donor tag sequence (excluding the N57 binding sequence) was codon optimised to increase the efficiency of mRNA translation (16) using the vector *Design Studio* tool from *VectorBuilder*, and a Kozak sequence was reestablished to facilitate the initiation of translation by ribosomes (17). The donor tag was purchased from *Integrated DNA Technologies* in the form of a custom gene in a vector flanked with HindIII and XhoI restriction sites to facilitate cloning (referred to as Sleeping ORANGE tag construct (SBpOR TagC) plasmids). **Figure 1** explains how endogenous tagging using the Sleeping ORANGE method works.

**Figure 1.**
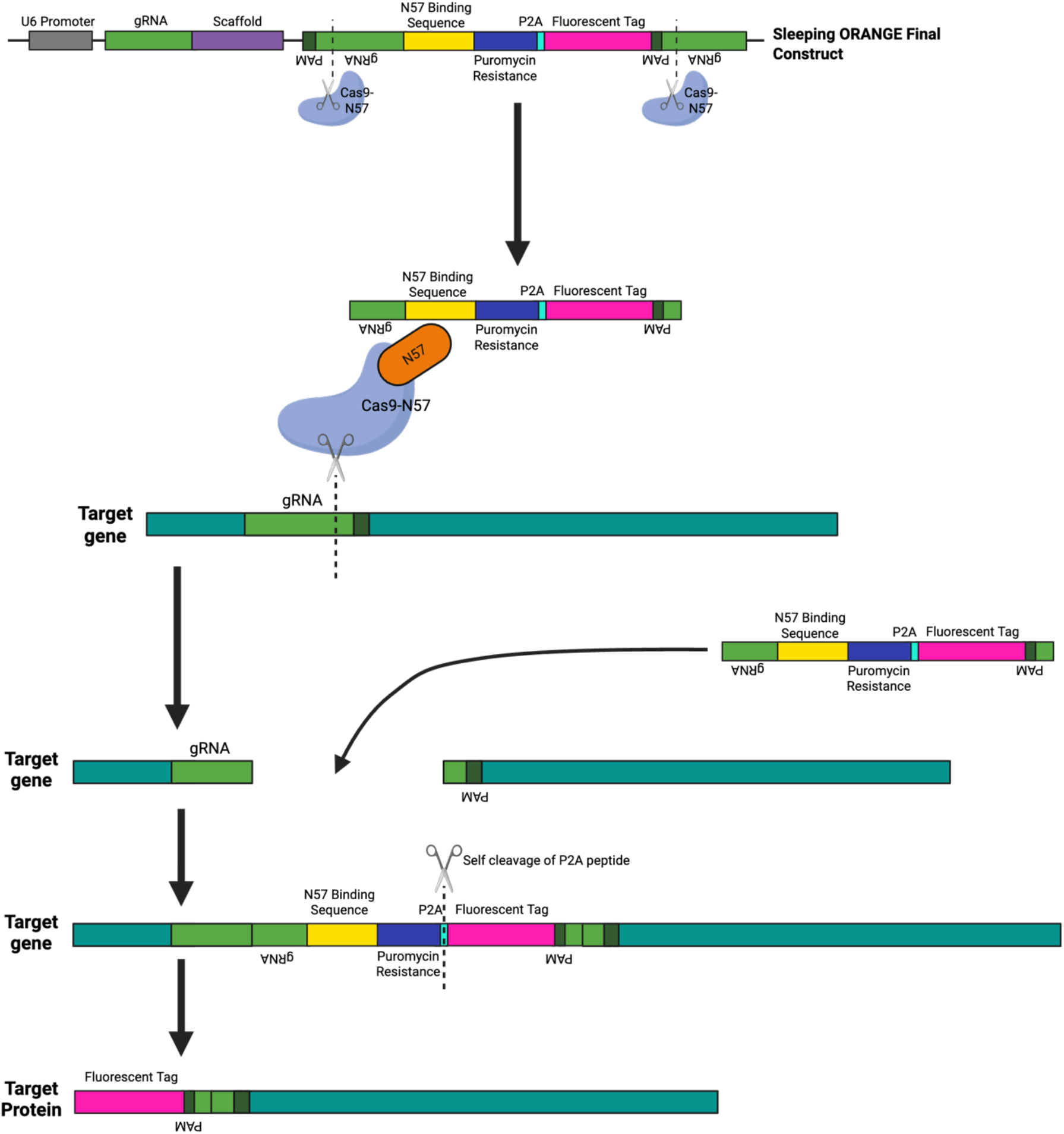
Endogenous epitope tagging through a novel CRISPR-Cas3 gene-editing technique, referred to as Sleeping ORANGE. Diagram illustrating how the Sleeping ORANGE technique uses CRISPR-Cas9 gene editing to facilitate the introduction of the fluorescent protein tag into the target gene. The Sleeping ORANGE final construct (SBpOR FC) encodes the sgRNA (under the control of a U6 promoter) and contains (in the following order) the N57 binding sequence which is composed of three DRs (the sequences the N57 domain of Sleeping Beauty binds to) along with IRs, a Puromycin resistance gene, a P2A peptide for self-cleavage and the donor sequence encoding the fluorescent tag. The SBpOR FC is co-transfected into cells alongside the pCAG-spCas9-N57 plasmid (#169920, *Addgene*). Within the cell, the Cas9-N57 fusion protein, guided by the sgRNA sequences, cleaves out the fluorescent tag sequence from the SBpOR FC as well as creating a double-stranded break at the target position in the target gene. The fluorescent tag-containing donor sequence can then be integrated into the genome by NHEJ without the need for sequence homology. The binding of Cas9-N57 to the N57 binding sequence in the donor sequence ensures that the donor sequence is brought to the insertion site to improve the editing efficiency. Note the ‘destruction’ of the gRNA-PAM sequences, preventing Cas9-N57 from inducing further double-stranded breaks in the DNA. When the modified gene is expressed, the P2A peptide encoded by the donor sequence is then self-cleaved, to separate the peptide translated from the N57 binding sequence and Puromycin resistance gene from the downstream fusion protein. Created with BioRender.com.

### Generating the pORANGE constructs

For each pORANGE construct, the gRNA sequences were cloned into the pORANGE Cloning template vector (referred to as the ‘empty vector’ (EV)) using type IIs restriction enzymes. The pORANGE Cloning template vector was a gift from Harold MacGillavry (Addgene plasmid #131471; http://n2t.net/addgene:131471; RRID: Addgene_131471). gRNAs were annealed and phosphorylated using T4 polynucleotide 5’-hydroxyl kinase (PNK; #EK0031, *Fisher Scientific*™). The pORANGE EV plasmid was digested using FastDigest BpiI (or BbsI) (#FD1014, *Fisher Scientific*™). The annealed gRNAs were ligated into the pORANGE EVs using T4 DNA Ligase (#EL0016, *Fisher Scientific*™), to generate the pORANGE intermediate constructs (pORANGE/pOR ICs).

The pORANGE ICs were then transformed into chemically competent DH5α *E. coli*, a single transformed colony picked for each construct and expanded, and plasmids extracted using the QIAprep® Spin Miniprep kit (#27106, *Qiagen*), following manufacturer instructions. Successful ligation of the gRNAs into the plasmids were confirmed by a double restriction digest with EcoRI and BbsI to confirm loss of BbsI restriction sites and plasmids with the correct digestion pattern were sent for Sanger sequencing (by *GENEWIZ*, *Azenta Life Sciences*) using the LKO.1 5’ primer (5’-GACTATCATATGCTTACCGT-3’) to confirm correct insertion of the gRNAs.

The pORANGE ICs as well as the pORANGE TagCs were then digested using FastDigest HindIII (#FD0504, *Fisher Scientific*™) and FastDigest XhoI (#FD0694, *Fisher Scientific*™) restriction enzymes, following manufacturer instructions. The digested donor tag sequences were then ligated into the respective digested pORANGE ICs using T4 DNA Ligase, generating the pORANGE final constructs (pORANGE/pOR FCs). pORANGE FCs were transformed into chemically competent DH5α *E. coli* and plasmids extracted as above. Successful ligation was confirmed by a triple restriction digest with HindIII, XhoI and EcoRI plasmids with the correct digestion pattern were sent for Sanger sequencing (by *GENEWIZ*, *Azenta Life Sciences*) using the LKO.1 5’ primer to confirm correct insertion of the donor tag.

### Generating the CYFIP1 Sleeping ORANGE construct

The CYFIP1 pORANGE IC and CYFIP1 Sleeping ORANGE TagC were digested using FastDigest HindIII and FastDigest XhoI restriction enzymes, following manufacturer instructions. The digested donor tag sequence was then ligated into the digested pORANGE IC using T4 DNA Ligase, generating the Sleeping ORANGE IC Step 2 (SBpOR IC S2). Sleeping ORANGE IC S2s were transformed into chemically competent DH5α *E. coli* and plasmids extracted as above. Successful ligation was confirmed by a triple restriction digest with HindIII, XhoI and EcoRI plasmids with the correct digestion pattern were sent for Sanger sequencing (by *GENEWIZ*, *Azenta Life Sciences*) using the LKO.1 5’ primer to confirm correct insertion of the donor tag sequence.

Sleeping ORANGE IC S2 was then digested with FastDigest BamHI (#FD0054, *Fisher Scientific*™) and FastDigest NotI (#FD0596, *Fisher Scientific*™), following manufacturer instructions, to cleave out the spCas9 sequence. A short “filler” sequence (CGTGACTGGGAAAACCCTGG) was designed containing a ‘useless’ sequence from the *E. coli LacZ* gene in *Benchling* and purchased from *Integrated DNA Technologies* (along with its complementary sequence) with sticky ends to facilitate cloning. This “filler” sequence was annealed and phosphorylated using T4 PNK, then ligated into the digested Sleeping ORANGE IC S2 using T4 DNA Ligase, generating the Sleeping ORANGE final construct (Sleeping ORANGE/SBpOR FC). SBpOR FCs were transformed into chemically competent DH5α *E. coli* and plasmids extracted as above. Successful ligation was confirmed by a triple restriction digest with BamHI, NotI and EcoRI and plasmids with the correct digestion pattern were sent for Sanger sequencing (by *GENEWIZ*, *Azenta Life Sciences*) using the LKO.1 5’ primer and the F1Ori primer (5’-CTCCTTTCGCTTTCTTCCT-3’) to confirm correct insertion of the donor tag sequence.

### Culture and transfection of HEK2G3 cells

Human embryonic kidney (HEK)-293 cells (#CRL-1573™, *American Type Culture Collection*) were thawed from frozen stocks stored in 10% dimethyl sulfoxide (#D2650-5X5ML, *Sigma-Aldrich*®) at −80°C. HEK293 cells were maintained in 10-cm-diameter Petri dishes (#P7612-360EA, *Sigma-Aldrich*®) with 10 mL of Dulbecco’s modified eagle medium/nutrient mixture F-12 with GlutaMAX (DMEM/F-12; #11524436, *Fisher Scientific*™) supplemented with 10% foetal bovine serum (FBS; #11550356, *Fisher Scientific*™), and incubated at 37°C, 5% CO2, 95% humidity. Cells were passaged at 70% confluency by incubation in 3 ml of pre-warmed (37°C) TrypLE™ (#10718463, *Fisher Scientific*™) for 5 min and TrypLE™ was inactivated by dilution in an equal volume of DMEM/F-12 + 10% FBS. 24 hours prior to transfection, 300,000 HEK293 cells were seeded into Nunc cell-culture treated 6-well plates (#10469282, *Fisher Scientific*™) in 3 mL of DMEM/F-12 + 10% FBS. The following day, HEK293 cells were transfected with plasmid DNA using Lipofectamine™ LTX Reagent (#15338100, *Fisher Scientific*™), following manufacturer instructions. For the Sleeping ORANGE system, HEK293 cells were co-transfected with the Sleeping ORANGE FC along with the pCAG-spCas9-N57 plasmid in a 2:1 ratio. The pCAG-spCas9-N57 plasmid was a gift from Zhili Rong (Addgene plasmid #169920; http://n2t.net/addgene:169920; RRID:Addgene_169920). Media was replaced 24 hours post-transfection.

### Puromycin selection for HEK2G3 cells edited using Sleeping ORANGE technique

7 days post-transfection with the CYFIP1 Sleeping ORANGE FC, HEK293 cells were treated with 1 μg/mL Puromycin (#P8833-10MG, *Sigma-Aldrich*®) in DMEM/F-12 + 10% FBS and Puromycin-containing media was replaced every 2-3 days for 14 days. This concentration of Puromycin was chosen based on our previous work (18).

### Single cell-derived colony isolation by picking

Following 14 days of 1 μg/mL Puromycin treatment, 100 and 1,000 cells were seeded into two separate 10-cm-diameter Petri dishes. Over the course of 1-2 weeks, these Petri dishes were monitored for the formation of discrete, likely arising from a single cell, colonies of cells. Colonies containing mNeonGreen-expressing cells were identified with the EVOS FLoid™ microscope (*Thermo Fisher Scientific*™) and a pipette was used to scrape and aspirate individual colonies and transport them into separate wells of a 24-well plate. The cells were then allowed to grow to confluency.

### Cell fixation, immunocytochemical staining and fluorescence imaging

For imaging of fixed cells, sterile 13-mm-diameter glass coverslips (#631-1578, *VWR*®) were seeded into 24-well plates and coated with 300 μL of 20 μg/mL poly-D-Lysine (PDL; #354210, *Corning*®) and incubated overnight at 37°C. 48 hours post transfection 30,000 HEK293 cells were seeded onto PDL-coated coverslips in 500 μL DMEM/F-12 + 10% FBS and incubated overnight. 24 hours later (72 hours post-transfection), HEK293 cells were fixed with 4% paraformaldehyde (PFA; pH = 7.4) (#P6148-1kg, *Sigma-Aldrich*®) for 20 minutes at room temperature. The following steps were all preceded with three 10-minute washes with 1x Phosphate-Buffered Saline (PBS) (#P4417-50TAB, *Sigma-Aldrich*®) and were performed at room temperature unless stated otherwise. Cells were permeabilized with 0.2% Triton X-100 (#T8787-100ML, *Sigma-Aldrich*®) in 1x PBS for 20 minutes, blocked with a solution of 0.1% Triton X-100 and 1% bovine serum albumin (BSA) (#A9647-50G, *Sigma-Aldrich*®) in 1x PBS for 1 hour, then sequentially incubated with 200 μL solutions of primary antibodies (2-3 hours at room temperature or overnight at 4°C) and secondary antibodies (1 hour at room temperature). For antibody specifics, see **Table 1**. All antibodies were diluted in blocking buffer. Nuclei were counterstained with 300 μL of 200 ng/mL 4′,6-diamidino-2-phenylindole (DAPI) (#D9542, *Sigma-Aldrich*®) for 15 minutes, then coverslips mounted onto glass microscope slides with ProLong™ Diamond Antifade Mountant (#15468070, *Fisher Scientific*™). Where needed, F-Actin was counterstained with 200 μL of 2 U/mL Alexa Fluor® 647-conjugated Phalloidin (#A22287, *Invitrogen™*) for 2 hours at room temperature prior to coverslip mounting. Microscope slides were imaged on the TCS SP8 confocal microscope (*Leica Microsystems*).

**Table 1:**
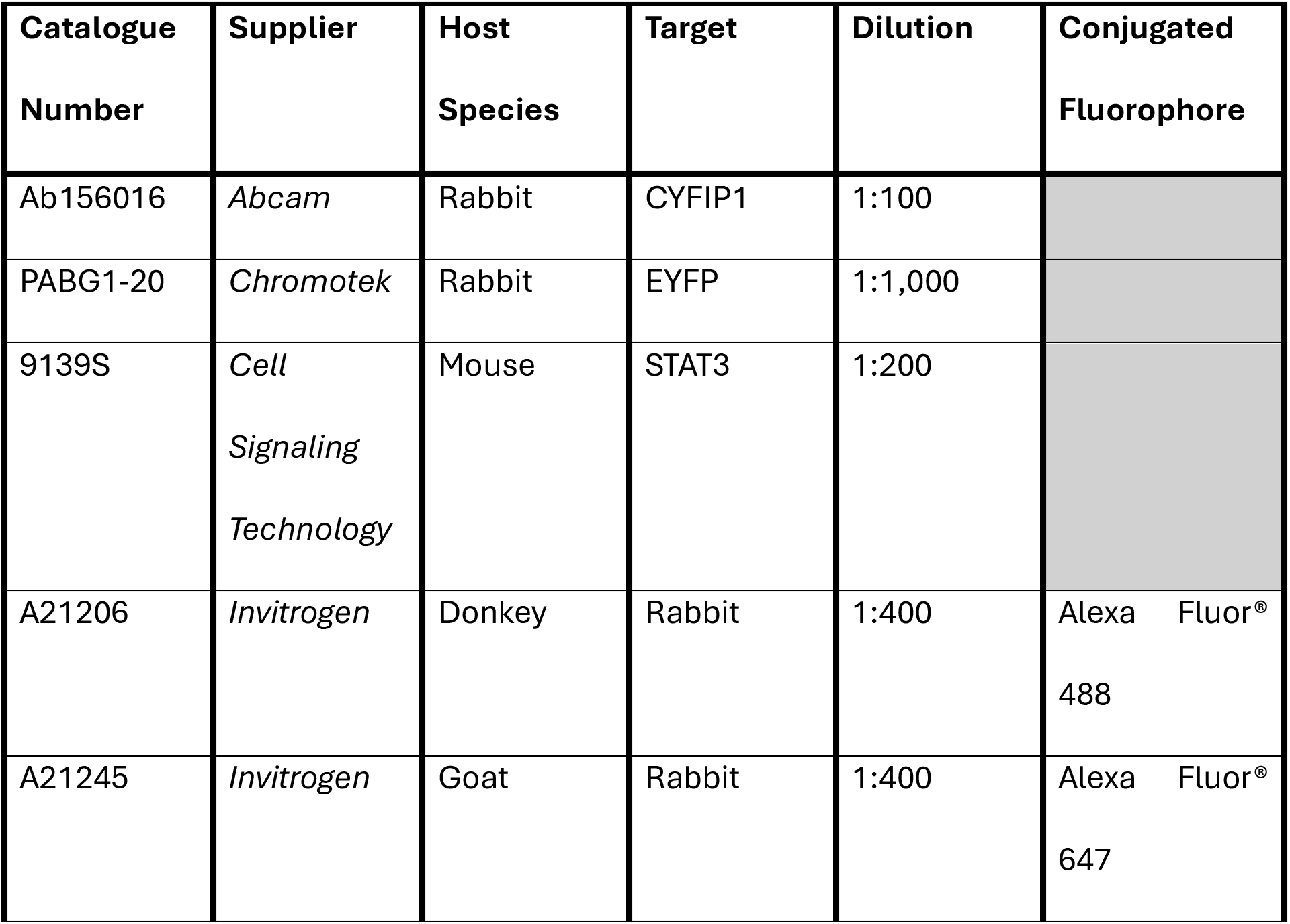
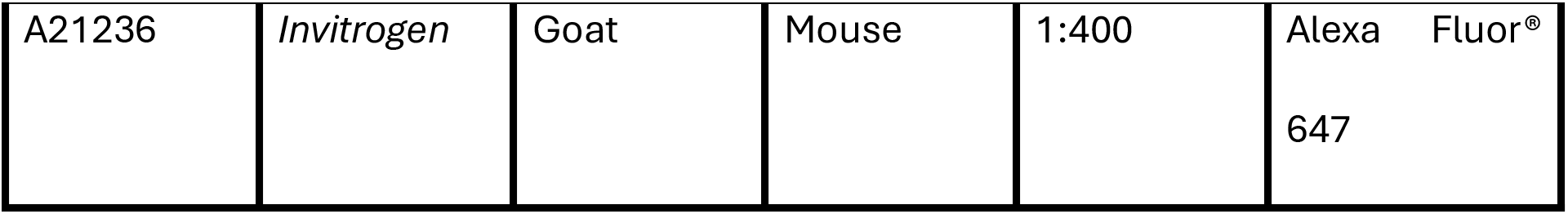
Antibodies used in this study.

To examine the proportion of mNeonGreen-expressing cells after ORANGE or Sleeping ORANGE gene-editing, cells were seeded into 6-well plates and transfected as described above, then left for 72 hours to allow for mNeonGreen expression. Nuclei were counterstained with by supplementation of 5 μg/mL Hoechst 33342 into the culture media for 30 minutes before live imaging on the DMi8 widefield microscope (*Leica Microsystems*). For higher-resolution live imaging of mNeonGreen localisation in the edited cells, isolated single cell-derived colonies were allowed to reach confluency before seeding onto PDL-coated glass-bottomed FluoroDish™ dishes (#FD3510-100, *World Precision Instruments*) (at a density of 150,000 cells/dish) and imaging on the TCS SP8 confocal microscope (*Leica Microsystems*). FluoroDish™ dishes were coated with 1 mL of 20 μg/mL PDL and incubated overnight as above.

### Flow cytometry

300,000 HEK293 cells were seeded into 6-well plates, then transfected the following day with either the CYFIP1 pORANGE FC or co-transfected with the CYFIP1 Sleeping ORANGE final construct and spCas9-N57 plasmid in a 2:1 ratio as described above, then examined for mNeonGreen fluorescence 72 hours post-transfection. As a positive control, CYFIP1 Sleeping ORANGE FC and pCAG-spCas9-N57-transfected cells that had undergone 2 weeks of Puromycin selection were seeded into a separate well. Cells were then harvested by detachment using TrypLE™, collected into a centrifuge tube with 2% FBS in 1x PBS (#14190144, *Fisher Scientific*™), then pelleted (200xg for 5 minutes). Cell pellets were resuspended in 1 mL of 2% FBS in 1x PBS and transferred into FACS tubes (#MTCT90051S, *Sigma-Aldrich*®). mNeonGreen signal intensities were then measured with the FITC filter of the Accuri™ C6 Flow Cytometer (*BD Biosciences*).

### DNA extraction, polymerase chain reaction and gel electrophoresis

Isolated clonal cell lines were grown to confluency in 24-well plates and then harvested by detachment using TrypLE™, collected into 1.5-mL tubes and pelleted (200xg for 5 minutes). Genomic DNA was extracted from cell pellets using the PureLink™ Genomic DNA Mini Kit (#K182001, *Invitrogen™*) following manufacturer’s instructions. DNA concentration, quality and purity was determined with a NanoDrop™ ND-8000 spectrophotometer (*Thermo Fisher Scientific™*).

20 μL PCR reactions were set-up using final concentrations of 50 ng genomic DNA, 0.5 μM Forward Primer and Reverse Primer (*Integrated DNA Technologies*), 1x B1 Buffer, 1.875 mM MgCl_2_, 0.2 mM dNTPs, 0.5 units of HOT FIREPol® DNA Polymerase (#01-02-KIT-0000S, *Solis BioDyne*). Thermocycler conditions were 95°C for 15 minutes; 35 cycles of: 95°C for 20 seconds, 60°C for 30 seconds, 72°C for 2.5 minutes; 72°C for 15 minutes, 15°C hold. To determine correct insertion of mNeonGreen sequence into *CYFIP1*, a forward primer in the mNeonGreen sequence (5’-TGGTGCCGCTCTAAGAAGAC-3’) and a reverse primer in the *CYFIP1* exon 4 (5’ATTGAACAAGCCACCGTCCA-3’) were designed and purchased from *Integrated DNA Technologies*. As a DNA ‘loading’ control, a forward primer (5’-CAACTTTGCGACATAAATTTTGGGG-3’) and reverse primer (5’-GCTGTACTGGGCGTTGTACT-3’) in the *NKX2-2* gene were used.

PCR products were mixed with 6x orange DNA loading dye (#11551575, *Fisher Scientific™*) and run alongside 1kb and 100bp DNA ladders (#N0550S and #N0551S, *New England Biolabs*) on 1.5% agarose gels (#10776644, *Fisher Scientific™*) in 1x TAE buffer (#574797-1L, *Sigma-Aldrich*®) with 1x SYBR™ Safe DNA Gel Stain (#10328162, *Invitrogen™*) at 120V for 45 minutes. Gels were imaged on the *UVP GelMax Imager*.

## Results

### Generating pORANGE constructs to endogenously tag genes of interest with fluorescent proteins

Using the pORANGE ‘empty vector’ (EV) backbone plasmid, we designed pORANGE constructs to endogenously tag three proteins of interest with three distinct fluorescent tags: CYFIP1 with the green fluorescent protein, mNeonGreen; JAKMIP1 with the red fluorescent protein, mScarlet; and STAT3 with the yellow fluorescent protein, mGold (**Figure 2**). To confirm that the constructs were correctly generated, restriction enzyme diagnostic digests and Sanger sequencing were performed. HEK293 cells were transfected with each pORANGE FC and the fluorescent signals emitted from gene-edited cells imaged. To confirm that the observed tagged fluorescent protein fusions localise to expected cellular compartments (and that the inclusion of the fluorescent protein did not affect the subcellular localisation of the tagged protein), transfected HEK293 cells were either stained with an antibody against the endogenously expressed tagged protein of interest (either CYFIP1 or STAT3), or in the case of JAKMIP1 where no validated antibody was available (at least for fluorescence microscopy), HEK293 cells were co-transfected with a previously published plasmid encoding a JAKMIP1-YFP fusion construct. We observed that the CYFIP1-mNeonGreen, JAKMIP1-mScarlet and STAT3-mGold fusion proteins appear to co-localise with the associated antibody staining or overexpressed JAKMIP-YFP construct (**Figure 2**). This suggests that the pORANGE plasmids correctly inserted the sequences encoding the respective fluorescent proteins into the desired genes. However, both CYFIP1-mNeonGreen and JAKMIP1-mScarlet also appear to unexpectedly localise to the nucleus (**Figure 2**), suggesting that perhaps there may be some off-target effects or that the mNeonGreen and mScarlet tags could be cleaved off, passively diffusing into the nucleus.

**Figure 2.**
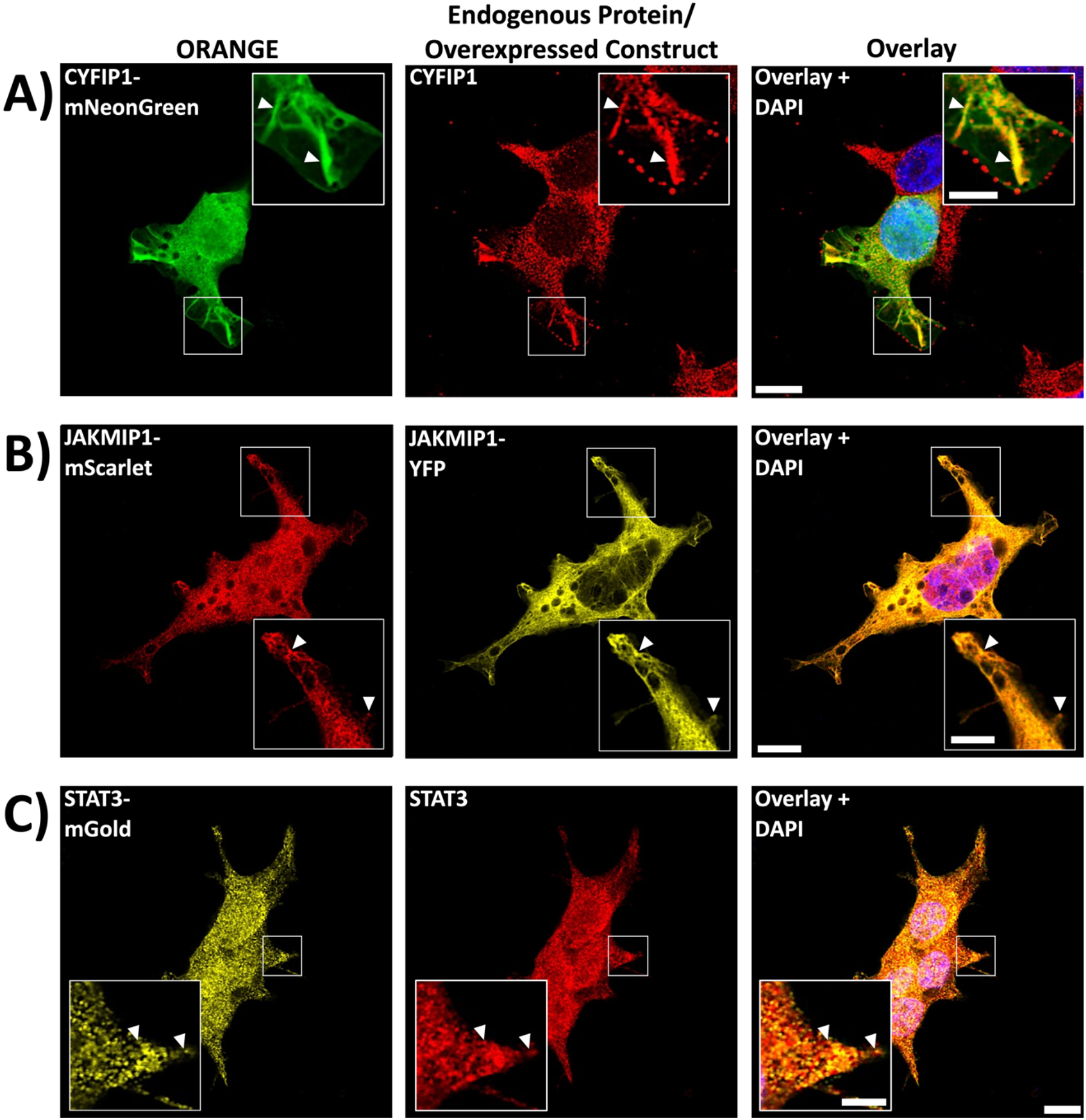
Localisation of CYFIP1-mNeonGreen, JAKMIP1-mScarlet and STAT3-mGold in HEK233 cells. HEK293 cells were transfected with the pORANGE FCs and fixed with 4% PFA 72 hours post-transfection. **A)** HEK293 cells transfected with the CYFIP1 pORANGE FC (to tag endogenous CYFIP1 with mNeonGreen) and stained with an antibody against CYFIP1 (in red). **B)** HEK293 cells co-transfected with the JAKMIP1 pORANGE FC (to tag endogenous JAKMIP1 with mScarlet) and the pCAβ-JAKMIP1-YFP plasmid. **C)** HEK293 cells transfected with the STAT3 pORANGE FC (to tag endogenous STAT3 with mGold) and stained with an antibody against STAT3 (in red). To boost the mGold signal, cells were co-stained with an anti-YFP antibody as the mGold signal alone was too faint following fixation. For all conditions, nuclei were counterstained with DAPI, and cells imaged on the *Leica* TCS SP8 inverted scanning confocal microscope. Arrowheads indicate examples of co-localisation. Scale bar = 10 μm. Inset scale bar = 2 μm.

Although the pORANGE constructs appear to be correctly tagging their respective proteins of interest, the tagging efficiency of these constructs is very low. An example demonstrating the poor editing efficiency of the JAKMIP1 pORANGE FC can be found in **Supplementary Figure 1**. Note that similar editing efficiencies were observed with both the CYFIP1 and STAT3 pORANGE FCs (data not shown here). In case the modified fusion proteins were simply lowly expressed, therefore making it difficult to identify gene-edited cells by fluorescence microscopy, we performed fluorescence-activated cell sorting as this would be more sensitive at detecting weakly fluorescent cells. This revealed that the percentage of cells exhibiting fluorescence for each construct was less than 2% (data not shown here), which agrees with the low number of gene-edited cells observed by eye through microscopy.

### Creating a novel method for endogenous epitope tagging based on ORANGE, referred to as Sleeping ORANGE

To try and improve the editing efficiency associated with endogenous tagging by ORANGE, we decided to combine the ORANGE technique with another technique that has been demonstrated to improve the efficiency of HITI (15). This technique makes use of the Sleeping Beauty transposase, more specifically the DNA binding PAI domain of the transposase (encoded within the 57 N-terminal amino acids of the transposase; referred to as N57 by (15)), and uses a fusion protein composed of this DNA binding domain attached to Cas9 (referred to as Cas9-N57). Fusing this domain to Cas9 combines the editing precision of the CRISPR-Cas9 system with the DNA-binding capabilities of the Sleeping Beauty transposase (15). Thus, when the N57-binding sequence is added to a donor tag, this system allows the donor tag sequence to be tethered to the fusion Cas9-N57 protein so that it is in close proximity to the target locus, facilitating integration.

A Sleeping ORANGE FC was then created using the same pORANGE intermediate construct (IC) for *CYFIP1* to tag endogenous CYFIP1 with mNeonGreen. To confirm that the construct was correctly generated, restriction enzyme diagnostic digests and Sanger sequencing were performed. HEK293 cells were co-transfected with the CYFIP1 Sleeping ORANGE FC and the pCAG-spCas9-N57 plasmid. Puromycin selection was used to generate an enriched population of successfully transfected and edited HEK293 cells, and single cell-derived colonies were isolated. To confirm that the observed mNeonGreen signals localise as expected based on existing literature, isolated cell populations of CYFIP1-mNeonGreen HEK293 cells were stained with an antibody against CYFIP1. We observed that the CYFIP1-mNeonGreen appeared to co-localise with the CYFIP1 staining (**Figure 3A**). Moreover, CYFIP1 is known to associate with F-Actin-rich structures and regions (19), and we observed that CYFIP1-mNeonGreen appears to localise to the membrane periphery and that it is often concentrated in cellular projections, regions which are likely to be rich in F-Actin (**Figure 3B-E**). For example, CYFIP1-mNeonGreen appears to accumulate at the distal end of cellular processes (**Figure 3B and C**), lamellipodia-like structures (**Figure 3D**), and potentially membrane-substrate contact sites (**Figure 3E**). Together, these observations suggest that the mNeonGreen is likely correctly tagging CYFIP1. However, similar to the CYFIP1-mNeonGreen localisation in HEK293 cells tagged using the pORANGE construct, CYFIP1-mNeonGreen appears to mis-localise to the nucleus in HEK293 cells tagged with mNeonGreen using the Sleeping ORANGE technique (**Figure 3A**).

**Figure 3.**
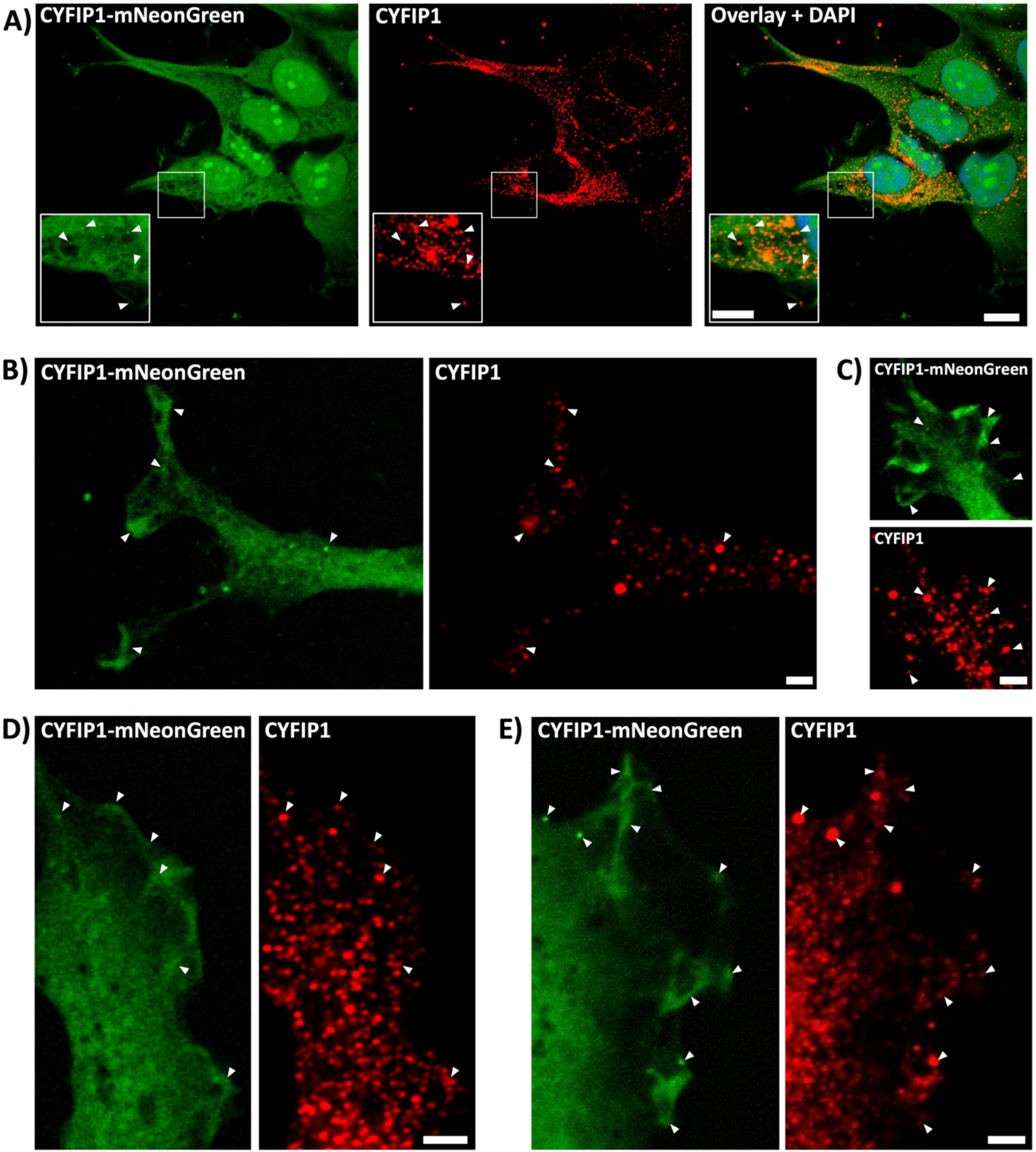
Localisation of CYFIP1-mNeonGreen in HEK233 cells following Sleeping ORANGE gene editing. **A)** HEK293 cells were co-transfected with the CYFIP1 SBpOR FC (to tag endogenous CYFIP1 with mNeonGreen) and pCAG-spCas9-N57, and successfully transfected cells selected for using Puromycin treatment for 14 days. Selected cells were isolated as single cell colonies, seeded onto coverslips, the fixed with 4% PFA. Cells were stained with an antibody against CYFIP1 (in red). Nuclei were counterstained with DAPI, and cells imaged on the *Leica* TCS SP8 inverted scanning confocal microscope. Arrowheads indicate co-localisation of CYFIP1-mNeonGreen and CYFIP1 antibody staining. Scale bar = 10 μm. Inset scale bar = 5 μm. **(B-E)** Images provide examples of CYFIP1-mNeonGreen localising at **B-C)** the distal end of cellular processes; **D)** lamellipodia-like structures; and **E)** likely membrane-substrate contact sites. Arrowheads indicate co-localisation of CYFIP1-mNeonGreen and endogenous CYFIP1 antibody staining. Scale bar = 2 μm.

### Sleeping ORANGE improves gene editing efficiency as compared to ORANGE

To compare the editing efficiency of the Sleeping ORANGE technique with the original ORANGE technique, HEK293 cells were transfected with either the CYFIP1 pORANGE FC, or co-transfected with the CYFIP1 Sleeping ORANGE FC and pCAG-spCas9-N57 plasmids, then imaged after 72 hours (**Figure 4A**). The number of mNeonGreen-positive (mNeonGreen^+^) cells and total number of cells were counted per field, and these values were used to calculate the percentage of mNeonGreen+ cells per field (**Figure 4B**). Although low, the percentage of mNeonGreen-expressing cells was roughly 12.85-fold higher when cells were edited with the Sleeping ORANGE technique (average of 1.518% of the population) compared to the ORANGE technique (average of 0.1181% of the population) (**Figure 4B**). This increase in edited cells is even more apparent when examining the proportion of fields sampled that contain mNeonGreen-expressing cells. Only 1 of the 22 analysed images (4.55%) contained mNeonGreen^+^ cells following ORANGE editing, whereas up to 11 of the 20 analysed images (55%) contained mNeonGreen^+^ cells after Sleeping ORANGE editing (**Figure 4C**). Flow cytometry was then used to confirm that populations of cells edited with the CYFIP1 Sleeping ORANGE FC showed higher mNeonGreen fluorescence compared to the wells of cells edited by the standard ORANGE technique (approximately 50% increase in mNeonGreen brightness) (**Figure 4D**), likely reflecting an increase in the number of gene-edited and therefore, mNeonGreen-expressing cells.

**Figure 4.**
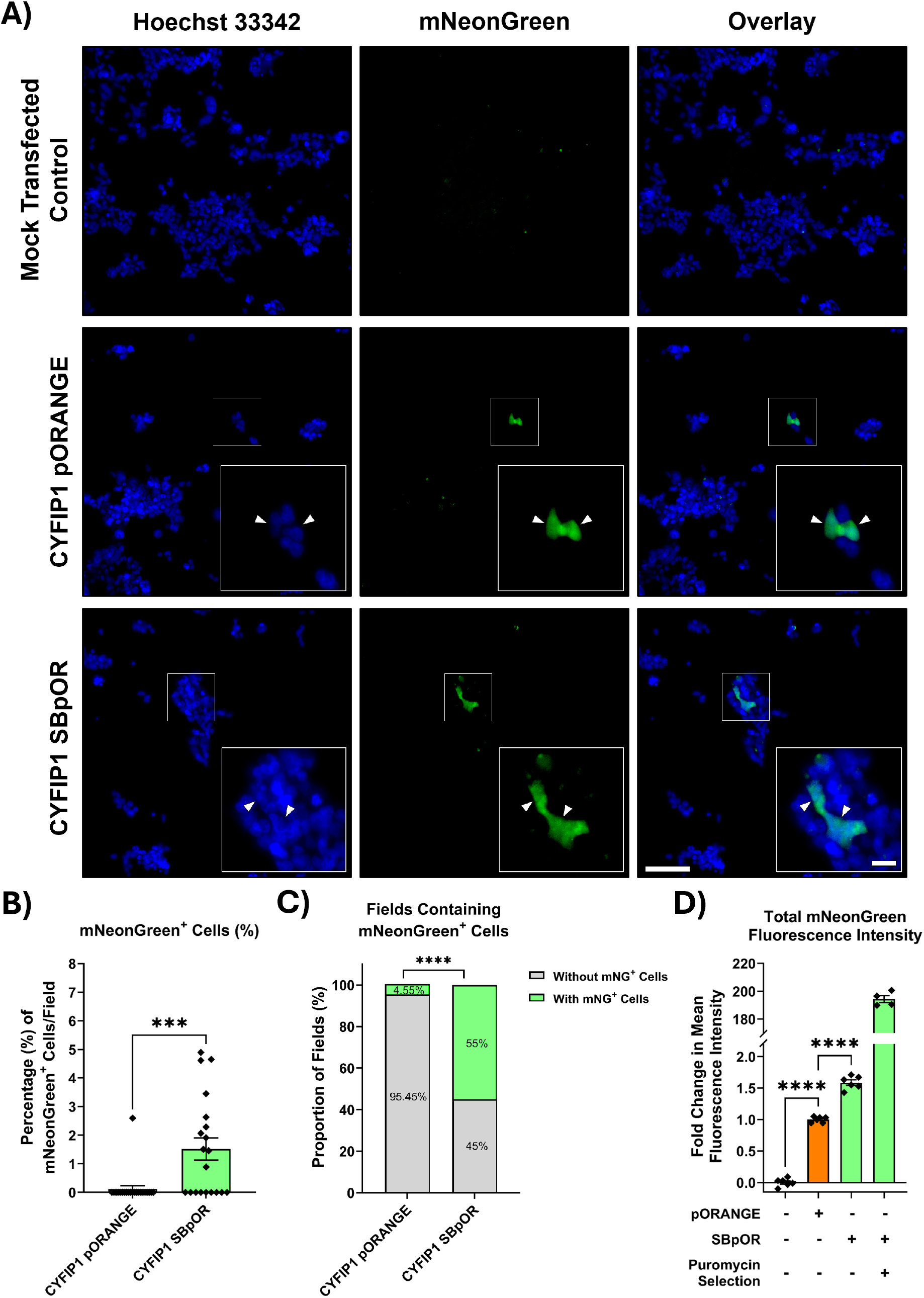
The Sleeping ORANGE system exhibits greater gene-editing efficiency compared to the ORANGE method. **A)** Live HEK293 cells examined for mNeonGreen fluorescence following ORANGE or Sleeping ORANGE gene-editing. HEK293 cells were seeded into 6-well plates at 300,000 cells/well, transfected with either the CYFIP1 pORANGE FC plasmid or a 2:1 mixture of the CYFIP1 SBpOR FC and pCAG-spCas9-N57 plasmids, and examined for mNeonGreen fluorescence after 72 hours. As a negative control, mock-transfected cells were seeded into a separate well. Nuclei were counterstained with Hoechst 33342 prior to imaging. White arrowheads indicate mNeonGreen-expressing cells. Scale bar = 100 mm. Inset scale bar = 20 mm. **B** and **C)**. Approximately 20-22 fields of view across four independently transfected wells of cells were chosen at random for analysis from cells depicted in A) (a total of 4-5 images per individually transfected well). **B)** The percentage of mNeonGreen^+^ cells per field were calculated for cells edited with the CYFIP1 pORANGE FC and SBpOR FC plasmids. Values displayed as mean ± SEM. Statistical significance of mean difference was determined using unpaired T-Test; ***P < 1x10^-3^. **C)** The proportion of fields containing mNeonGreen^+^ (abbreviated mNG^+^ in legend) cells per field were calculated for cells edited with the CYFIP1 pORANGE FC and SBpOR FC plasmids. Values displayed as mean ± SEM. Statistical significance of the difference in proportions was determined using Fisher’s exact test; ****P < 1x10^-4^. **D)** Fold change in mean mNeonGreen signal intensities measured with the FITC filter of a Accuri™ C6 Flow Cytometer (BD Biosciences) from six individually transfected wells of cells from part A). Four wells of Puromycin selected CYFIP1 SBpOR cells were used as a positive control. Values normalised to CYFIP1 pOR FC-transfected cells and displayed as mean ± SEM of three independently transfected wells, with the exception of the Puromycin selected CYFIP1 SBpOR transfected population of cells, data of which comes from only four wells. Statistical significance was determined using one-way ANOVA and P-values were adjusted by Tukey’s Honest Significant Difference test; ****P < 1x10^-4^.

### Variable mNeonGreen localisation across Sleeping ORANGE clones and screening for *mNeonGreen* integration by PCR

We noticed that the localisation of the mNeonGreen signal varies across and within the isolated cell populations can differ and drastically changes following fixation. **Figure 5** illustrates some of the different localisation patterns of mNeonGreen observed in the live SBpOR FC-transfected HEK293 cells. Importantly, although not quantified, the mNeonGreen signal seems to differ greatly in brightness between these isolated cell populations as well, suggesting potential differences in zygosity of copy number of *mNeonGreen* sequences inserted (**Figure 5**).

**Figure 5.**
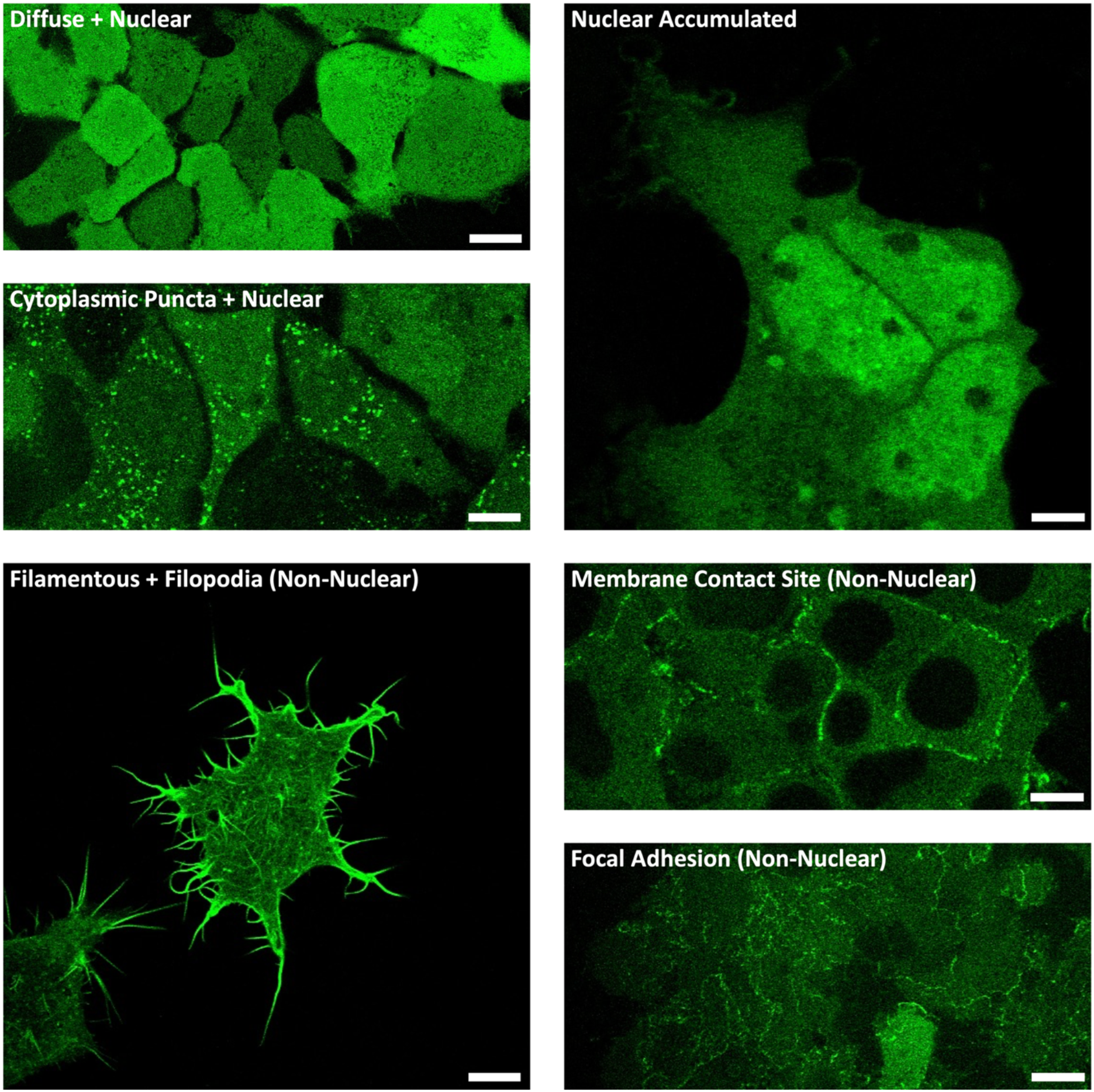
CYFIP1-mNeonGreen shows distinct localisations amongst different isolated populations of live HEK233 cells. HEK293 cells were co-transfected with the CYFIP1 SBpOR FC (to tag endogenous CYFIP1 with mNeonGreen) and pCAG-spCas9-N57, and successfully transfected cells selected for using Puromycin treatment for 14 days. Selected cells were isolated as single cell colonies and cells imaged on the *Leica* TCS SP8 inverted scanning confocal microscope. Scale bar = 10 μm.

Given these different mNeonGreen localisation patterns, we aimed to determine whether *mNeonGreen* was correctly integrated at the target site within *CYFIP1* in any of the isolated clones. Genomic DNA was extracted from clonal isolated cell populations and PCRs performed using a forward primer against part of the mNeonGreen sequence, and a reverse primer against a sequence in exon 4 of *CYFIP1* (**Figure 6A**). Therefore, PCR amplicons can only be produced if the mNeonGreen is inserted into *CYFIP1* at the desired site. **Figure 6B** shows that clonal lines “N2”, “N5” and “35” all show correct insertion of mNeonGreen into *CYFIP1*, whereas clonal lines “5” and “15” do not. Of a total 59 mNeonGreen-expressing clones screened, we found that the *mNeonGreen* sequence was inserted at the target locus in *CYFIP1* for three clones (**Figure 6C**).

**Figure 6.**
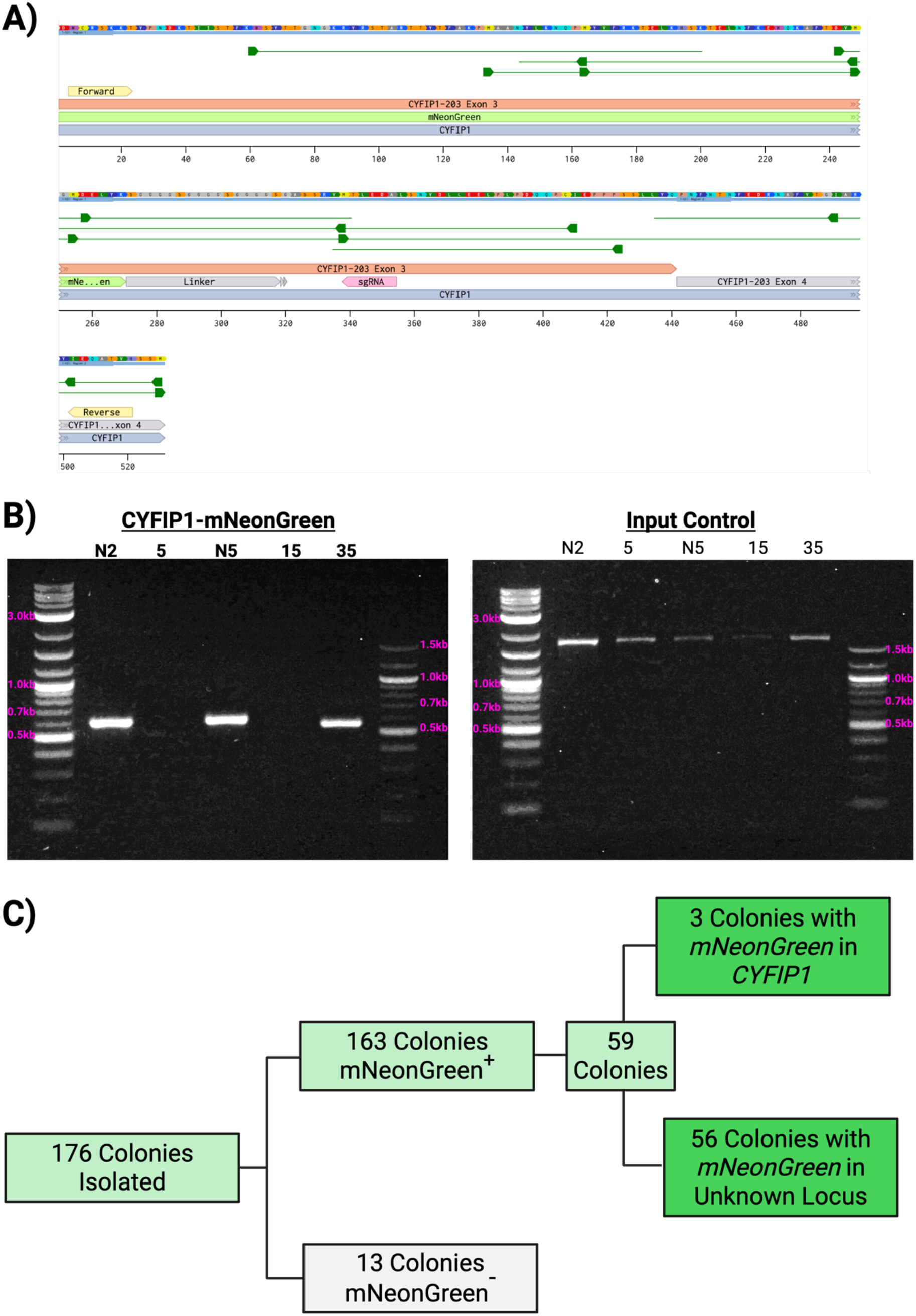
PCR screening identifies mNeonGreen integration at the CYFIP1 locus. **A)** Schematic of the PCR strategy used to detect correct integration of *mNeonGreen* into the *CYFIP1* locus. A forward primer anneals within the *mNeonGreen* sequence, while a reverse primer anneals within *CYFIP1* exon 4 downstream of the insertion site, enabling amplification only upon correct genomic integration. PCRs were performed on genomic DNA isolated from individual clonal cell lines. **B)** Representative agarose gel showing PCR products from selected clonal lines (N2, 5, N5, 44, and 35) using the primers described in A). Primers targeting the *NKX2-2* gene were used as a genomic DNA input control. **C)** Summary of clonal isolation and screening results. A total of 176 clonal lines were isolated following Puromycin selection, of which 163 were mNeonGreen positive and 13 were negative. 59 mNeonGreen⁺ clones were taken forward at random and screened by PCR, 3 of which showed correct integration at the CYFIP1 locus, while 56 exhibited integration at an unknown locus.

Importantly, clonal lines with *mNeonGreen* integrated into *CYFIP1* (clones #35, #N2 and #N5) display close association or the mNeonGreen signal with F-Actin, with the mNeonGreen being more enriched at processes or regions at the cell edges with more prominent F-Actin structures, and do not show the unexpected nuclear or nucleolar mNeonGreen accumulation (**Figure 7**). Furthermore, clonal lines where *mNeonGreen* is integrated into an unknown off-target locus do not show the same degree of mNeonGreen association with F-Actin, displaying increased mNeonGreen nuclear accumulation (clone #5) or more diffuse nucleocytoplasmic mNeonGreen patterning (clone #44) (**Figure 7**).

**Figure 7.**
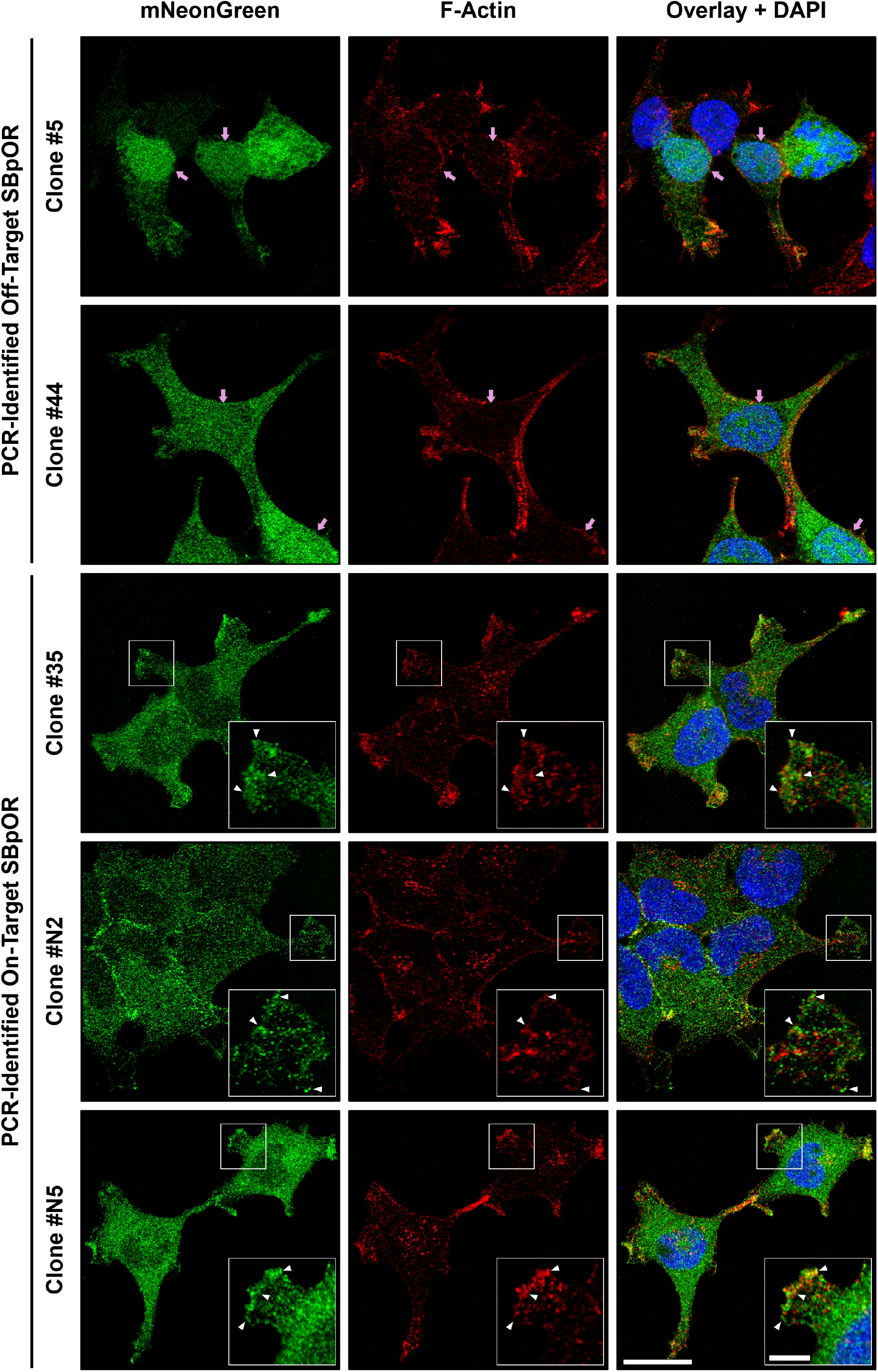
Close association of mNeonGreen-tagged CYFIP1 with F-Actin. **A)** HEK293 cells were co-transfected with the CYFIP1 SBpOR FC (to tag endogenous CYFIP1 with mNeonGreen) and pCAG-spCas9-N57, and successfully transfected cells selected for using Puromycin treatment for 14 days. Selected cells were isolated as single cell-derived colonies, expanded, screened for *mNeonGreen* integration into the *CYFIP1* gene by PCR, and then selected clones were seeded onto coverslips and fixed with 4% PFA. Clones #5 and #44 are examples of cell populations with *mNeonGreen* insertion at an unknown off-target locus. Clones #35, #N2 and #N5 are confirmed to have mNeonGreen insertion within *CYFIP1*. Nuclei and F-Actin were counterstained with DAPI and Phalloidin, and cells imaged on the *Leica* TCS SP8 inverted scanning confocal microscope. Pink arrows indicate nuclear localisation of mNeonGreen observed in the off-target clones. White arrowheads indicate close association or co-localisation of CYFIP1-mNeonGreen and F-Actin. Scale bar = 10 μm. Inset scale bar = 5 μm.

## Discussion

### Increased gene-editing efficiency with the Sleeping ORANGE technique

Visualising the subcellular localisation of proteins is essential for understanding cellular processes and disease mechanisms. Endogenous protein tagging makes use of fluorescent proteins or various other tags to label proteins of interest in their natural state, enabling them to be easily imaged even when no validated antibodies are available. The CRISPR-Cas9-based ORANGE method for endogenous tagging has previously been demonstrated to effectively tag an array of proteins with various reporters in neuronal cultures which are difficult to genetically edit (2). Therefore, we used ORANGE to endogenously tag three proteins, CYFIP1, JAKMIP1 and STAT3 with three fluorescent markers, mNeonGreen, mScarlet and mGold. We demonstrate that the fluorescent signals for all three ORANGE-tagged proteins show the expected subcellular localisations based on previous work and literature, and moreover, that the fluorescent signals from the fusion proteins co-localise with antibody staining against the target tagged protein (or overexpressed fluorescently tagged fusion protein in the case of JAKMIP1). Together, these results suggest that the sequences encoding the fluorescent proteins were likely inserted into the target genes of interest and did not affect the structure of the translated proteins.

Despite the promising localisations of the tagged proteins using the ORANGE method, we observed that the editing efficiency of the endogenous tagging was very low in HEK293 cells, a cell line often reported as being easy to transfect and which are commonly used for gene-editing studies (20). Therefore, in order to try and improve the editing efficiency of the ORANGE method, we combined the ORANGE method with a previously published technique shown to improve HITI-based CRISPR-Cas9 gene editing (15). Ma *et al.* hypothesised that fusing the DNA-binding sequence of the Sleeping Beauty transposon to Cas9 (referred to as Cas9-N57), could be used to bring the donor DNA template close to the Cas9 cleavage site, thereby encouraging DNA integration (15). They then demonstrated that correct integration of a sequence encoding GFP within the 3’ untranslated region (3’ UTR) of the *Actin Beta*(*ACTB*) gene was improved by over 20% compared to with Cas9 on its own (15), suggesting that this Cas9-N57 fusion protein improves the editing efficiency of HITI-dependent CRISPR-Cas9 gene editing. We therefore designed and generated a modified pORANGE FC that included the N57-binding sequence, which we termed Sleeping ORANGE, to better endogenously tag CYFIP1 with mNeonGreen. We co-transfected the CYFIP1 Sleeping ORANGE FC along with the pCAG-spCas9-N57 plasmid into HEK293 cells, then used a combination of Puromycin selection and single cell picking to isolate successfully transfected cell populations. We observed that transfected HEK293 cells expressed mNeonGreen, and following staining with an antibody against CYFIP1, found that the tagged CYFIP1-mNeonGreen co-localised with CYFIP1 antibody staining. Furthermore, the CYFIP1-mNeonGreen appeared to associate with expectedly F-Actin-rich structures, such as lamellipodia-like structures, although further staining for F-Actin is required to confirm this. This suggests that we were successfully able to tag endogenously expressed CYFIP1 with mNeonGreen using the Sleeping ORANGE method.

Importantly, we observed that the editing efficiency using the Sleeping ORANGE technique was higher than using the standard ORANGE technique. The percentage of mNeonGreen-expressing cells and fields containing mNeonGreen-expressing cells was higher, and flow cytometry showed greater mNeonGreen signal intensities in populations of cells transfected with the CYFIP1 Sleeping ORANGE FC and pCAG-spCas9-N57 plasmid compared to cells transfected with the CYFIP1 pORANGE FC. Together, these finding suggest that the Sleeping ORANGE method appears to be more effective for endogenous tagging than the ORANGE method on which it is based. However, whether the addition of the Sleeping Beauty DNA binding sequence and use of Cas9-N57 was the reason for the improved editing efficiency is unknown, as without directly investigating this, we have no direct evidence that Cas9-N57 recruited the donor tag to the cleavage site as expected. Instead, it may have been one of the other alterations made to the Sleeping ORANGE donor tag that improved the editing efficiency, such as the codon optimisation or reconstruction of the Kozak sequence. Importantly though, this technique has improved on the ORANGE technique for endogenous protein tagging.

### Off-target gene-editing with the ORANGE and Sleeping ORANGE techniques

Although the incorporation of the Cas9-N57 protein and N57-binding sequence improved gene-editing efficiency, we noted that fluorescent proteins tended to be mis-localised in populations of both ORANGE and Sleeping ORANGE edited cells. For example, using the ORANGE technique, we observed both mNeonGreen (designed to tag CYFIP1) signal and mScarlet (designed to tag JAKMIP1) signals in the nucleus. Similarly, we also observed mNeonGreen signal in the nucleus, and more specifically, high mNeonGreen nucleolar accumulation when tagging CYFIP1 using the Sleeping ORANGE method. This does not fit well with the known localisation of CYFIP1 given that there is no evidence to show CYFIP1 in the nucleus, and although JAKMIP1 has been demonstrated to exist in the nucleus, it predominantly localises to the cytoplasm (21). There are several potential explanations for this unexpected nuclear signal. The first being that it is not uncommon for fluorescent proteins to be cleaved or ‘fall off’ from recombinant proteins. Once cleaved, these fluorescent proteins are small enough to passively diffuse from the cytosol into the nucleus through the nuclear pores (22). This could be confirmed by subcellular fractionation and immunoprecipitation experiments to analyse whether the fluorescent proteins in the nuclear compartment remain bound to their target tagged protein.

However, if the fluorescent tags are indeed tagging their target proteins and are not being cleaved off, it is possible that the insertion of the fluorescent proteins could be altering the structure and folding of the tagged CYFIP1 and JAKMIP1 proteins, causing them to mis-localise to the nucleus. This could be assessed by examining cellular functions or pathways these proteins are known to be involved in to check for any irregularities that would indicate reduced or altered functionality, or by replacing the current fluorescent tags with short peptide tags that are less likely to affect the folding, function and localisation of CYFIP1 or JAKMIP1 (23). Considering that both CYFIP1 and JAKMIP1 have been successfully tagged with fluorescent proteins in previous studies without reports of mis-localisation (24–26), we do not expect that this is a major contributor to the unexpected nuclear or nucleolar accumulation of ORANGE-/Sleeping ORANGE-tagged proteins.

The final explanation for this unexpected nuclear localisation is that it most likely arises from incorrect integration of the fluorescent protein sequences into the genome. CRISPR-Cas9 gene-editing can create indels, leading to frame-shift mutations disrupting the coding sequences of these genes which could result the expression of the fluorescent protein with only a small, truncated region of the target protein that retains its ability to diffuse into the nucleus. Alternatively, the design of the gRNA could be suboptimal, causing the fluorescent protein sequences to be inserted into different regions of the genome through off-target gene editing and we may instead be observing unintentional tagging of other, nuclear-localising proteins. This appears to be the case with the Sleeping ORANGE technique as “CYFIP1-mNeonGreen” can show distinct localisations in different isolated cell populations, and PCR screening confirms that the majority of these clones do not even have the *mNeonGreen* sequence in the *CYFIP1* gene. Follow-up imaging demonstrates that the clones where *mNeonGreen* is detectably integrated into the *CYFIP1* gene show a lack of nuclear/nucleolar mNeonGreen accumulation and much more expected F-Actin-associated localisation as well. Although more thorough analysis of the edited cells should be performed as there are still some concerns that should be addressed before any downstream functional experiments are undertaken with these cells; for example, the zygosity of the edit and whether an indel mutation was introduced downstream of the mNeonGreen following HITI-mediated integration.

It is important to note that although it may appear that the Sleeping ORANGE method creates more off-target effects than the ORANGE method, these off-target effects would likely be observed in cells edited with the ORANGE technique given that the same gRNA sequences were used for both the ORANGE and Sleeping ORANGE techniques. However, we expect that the reason we did not observe the range of different protein localisations with the ORANGE-edited cells is simply because that the editing efficiency of the ORANGE technique was so low that only a few mNeonGreen-expressing cells were observed. The increased editing efficiency of the Sleeping ORANGE technique proportionally creates a greater chance to observe off-target effects but therefore, also a greater chance for on-target effects. We isolated three ‘on-target’ clones from screening 59 (an approximate on-target gene-editing success rate of ∼5% following Puromycin selection), which would be impractical with the base ORANGE technique (only one field from four independently transfected wells of cells contained a cluster of mNeonGreen-expressing cells). To reduce off-target gene-editing in future experiments, optimisation of gRNA choice and validation of gRNA efficiency is important as although the Sleeping ORANGE technique improves gene-editing efficiency, the binding kinetics of the specific gRNA and locations of the double-stranded breaks are the crucial denominators of on-/off-target gene-editing.

### Considerations for using Sleeping ORANGE potential improvements for future variants

It is important to note that the advantage of the ORANGE and Sleeping ORANGE techniques is that they are based on the HITI DNA repair pathway, and as such, enable endogenous tagging in post-mitotic cells (27). In our study, we have used the HEK293 cell line to test the efficiency of the Sleeping ORANGE technique. HEK293 cells were chosen for this as they are highly proliferative and typically amenable to CRISPR-based editing strategies. However, as HEK293 cells are not post-miotic cells, to evaluate the full potential of the Sleeping ORANGE technique, the technique should be tested using post-mitotic cells such as neurons. This would provide better insight into the true benefits and applicability of the Sleeping ORANGE technique.

The Sleeping ORANGE approach does have some potential further limitations due to the addition of the N57-binding sequence. To avoid making a new fusion protein containing not only the fluorescent protein tag but also the Puromycin resistance gene product and whatever peptide is encoded by the N57-binding sequence, we designed a polycistronic approach using a P2A ‘self-cleaving’ peptide sequence to separate these components from the rest of CYFIP1. We also identified a gRNA very close to the start of the first coding exon in *CYFIP1* to facilitate this, because any amino acids upstream of the gRNA would be removed when the P2A peptide sequence is skipped during translation after insertion of the donor tag into the genome. The loss of these amino acids causes two potential problems: the first being that if these N-terminal amino acids are important for folding or function of a given protein, losing them may result in a reduced or non-functional protein; the second being that this places strict limitations on gRNA design. This could be a problem for proteins with crucial functional domains at the N-terminus. The Sleeping ORANGE system could be modified to tag proteins at the C-terminus, as this would simply require flipping of the order of sequences in the donor tag to place the Puromycin resistance gene and N57-binding sequence at the 3’ end of the donor tag downstream of the fluorescent protein and P2A peptide sequences, although this would need to be tested.

Additionally, there is some concern about the peptide encoded by the N57-binding sequence. Although we have drastically shortened the length of this sequence to only contain the most important IR/DR sequences, it is unknown what the function of this peptide is. In the best case, it is an inert peptide with no cellular function, but its presence may still alter some cellular function or signalling pathway. So far, it does not appear to be cytotoxic, as the Sleeping ORANGE-edited cells survive the selection process and can be expanded without issue. To circumvent this issue, future attempts to use the Sleeping ORANGE system could target this peptide for degradation, perhaps by using degron tags, adding sequences that contain proteasome-targeting signals or similar approaches. With another P2A peptide sequence between the N57-binding sequence and the Puromycin resistance gene, this would mean the only extra product existing within the cell is the fluorescent protein (hopefully fused to the protein of interest) and the product of the Puromycin resistance gene.

## Conclusion

Overall, we have developed an endogenous protein tagging method with enhanced gene-editing efficiency compared to the initial ORANGE technique. With further work to optimise gRNA design to ensure that fluorescent protein sequences are correctly integrated into their target gene and reduce off-target effects, an improved Sleeping ORANGE technique can be a useful tool for researchers to better study protein subcellular localisation and dynamics.

## Declarations

### Ethics approval and consent to participate

This study did not involve human participants or human data and therefore no ethics approval was required for this study. The HEK293 cells used in this study are commercially available from the *American Type Culture Collection*.

### Consent for publication

Not applicable.

### Availability of data and materials

The datasets used and/or analysed during the current study are available from the corresponding author on reasonable request.

### Author’s contributions

E-R.M: Design, generation and validation of ORANGE constructs; design, generation and validation of Sleeping ORANGE construct; figure generation; manuscript writing and editing

J.G.M: Design of Sleeping ORANGE construct; validation of Sleeping ORANGE construct; critical insight; manuscript editing

K.A.L: Flow cytometry; manuscript editing

M.A.R: Project supervision; critical insight; manuscript editing

A.O-A: Project supervision; funding attainment; critical insight; manuscript editing

All authors reviewed the manuscript. All authors read and approved the final manuscript.

## Acknowledgements

The research was carried out at the National Institute for Health and Care Research (NIHR) Exeter Biomedical Research Centre (BRC). The authors would like to thank Lizzy Sears for assisting with maintaining HEK293 cell stocks. We also gratefully acknowledge Liming Yang and Madie Eve for their insightful discussions during the development of this work. The authors would also like to thank the Hospital Saturday Fund (HSF) and the Bristol Japanese Culture Society (BJCS) for providing charitable funding that supported this research.

## List of abbreviations

CRISPR: Clustered Regularly Interspaced Short Palindromic Repeats
Cas9: CRISPR associated protein 9
HITI: Homology-independent targeted integration
HDR: Homology-directed repair
NHEJ: Non-homologous end joining
PAM: Protospacer adjacent motif
gRNA: Guide RNA
ORANGE: Open Resource for the Application of Neuronal Genome Editing
GFP: Green fluorescent protein
RFP: Red fluorescent protein
YFP: Yellow fluorescent protein
CYFIP1: Cytoplasmic FMR1-interacting protein 1
STAT3: Signal transducer and activator of transcription 3
JAKMIP1: Janus kinase and microtubule-interacting protein 1
EV: Empty vector
IC: Intermediate construct
FC: Final construct
pOR: pORANGE
SBpOR: Sleeping ORANGE
WT: Wild type

## References

1. Husser MC, Pham NP, Law C, Araujo FRB, Martin VJJ, Piekny A. Endogenous tagging using split mNeonGreen in human iPSCs for live imaging studies. Elife. 2023;12:1–28. doi:10.7554/eLife.92819

2. Willems J, de Jong APH, Scheefhals N, Mertens E, Catsburg LAE, Poorthuis RB, et al. Orange: A CRISPR/Cas9-based genome editing toolbox for epitope tagging of endogenous proteins in neurons. PLoS Biol. 2020;18(4). doi:10.1371/journal.pbio.3000665

3. Conic S, Desplancq D, Tora L, Weiss E. Electroporation of Labeled Antibodies to Visualize Endogenous Proteins and Posttranslational Modifications in Living Metazoan Cell Types. Bio Protoc. 2018;8(21). doi:10.21769/BioProtoc.3069

4. Moriya H. Quantitative nature of overexpression experiments. Molecular Biology of the Cell. 2015. doi:10.1091/mbc.E15-07-0512

5. Schwinn MK, Steffen LS, Zimmerman K, Wood K V., Machleidt T. A Simple and Scalable Strategy for Analysis of Endogenous Protein Dynamics. Sci Rep. 2020;10(1). doi:10.1038/s41598-020-65832-1

6. Bukhari H, Müller T. Endogenous Fluorescence Tagging by CRISPR. Trends in Cell Biology. 2019. doi:10.1016/j.tcb.2019.08.004

7. Mahen R, Koch B, Wachsmuth M, Politi AZ, Perez-Gonzalez A, Mergenthaler J, et al. Comparative assessment of fluorescent transgene methods for quantitative imaging in human cells. Mol Biol Cell. 2014;25(22). doi:10.1091/mbc.E14-06-1091

8. Makhija S, Brown D, Rudlaff RM, Doh JK, Bourke S, Wang Y, et al. Versatile Labeling and Detection of Endogenous Proteins Using Tag-Assisted Split Enzyme Complementation. ACS Chem Biol. 2021;16(4). doi:10.1021/acschembio.0c00925

9. Schwinn MK, Machleidt T, Zimmerman K, Eggers CT, Dixon AS, Hurst R, et al. CRISPR-Mediated Tagging of Endogenous Proteins with a Luminescent Peptide. ACS Chem Biol. 2018;13(2). doi:10.1021/acschembio.7b00549

10. Uemura T, Mori T, Kurihara T, Kawase S, Koike R, Satoga M, et al. Fluorescent protein tagging of endogenous protein in brain neurons using CRISPR/Cas9-mediated knock-in and in utero electroporation techniques. Sci Rep. 2016;6. doi:10.1038/srep35861

11. Serebrenik Y V., Sansbury SE, Kumar SS, Henao-Mejia J, Shalem O. Efficient and flexible tagging of endogenous genes by homology-independent intron targeting. Genome Res. 2019;29(8). doi:10.1101/gr.246413.118

12. Suzuki K, Izpisua Belmonte JC. In vivo genome editing via the HITI method as a tool for gene therapy. Journal of Human Genetics. 2018. doi:10.1038/s10038-017-0352-4

13. Lambert TJ. FPbase: a community-editable fluorescent protein database. Nat Methods. 2019 Apr 18;16(4):277–8. doi:10.1038/s41592-019-0352-8

14. Chen X, Zaro JL, Shen WC. Fusion protein linkers: Property, design and functionality. Advanced Drug Delivery Reviews. 2013. doi:10.1016/j.addr.2012.09.039

15. Ma S, Wang X, Hu Y, Lv J, Liu C, Liao K, et al. Enhancing site-specific DNA integration by a Cas9 nuclease fused with a DNA donor-binding domain. Nucleic Acids Res. 2020;48(18). doi:10.1093/nar/gkaa779

16. Sen A, Kargar K, Akgün E, Plnar MC. Codon optimization: A mathematical programing approach. Bioinformatics. 2020;36(13). doi:10.1093/bioinformatics/btaa248

17. Ambrosini C, Destefanis E, Kheir E, Broso F, Alessandrini F, Longhi S, et al. Translational enhancement by base editing of the Kozak sequence rescues haploinsufficiency. Nucleic Acids Res. 2022;50(18). doi:10.1093/nar/gkac799

18. Harlow CE, Gandawijaya J, Bamford RA, Martin ER, Wood AR, van der Most PJ, et al. Identification and single-base gene-editing functional validation of a cis-EPO variant as a genetic predictor for EPO-increasing therapies. Am J Hum Genet. 2022;109(9). doi:10.1016/j.ajhg.2022.08.004

19. DeRubeis S, Pasciuto E, Li KW, Fernández E, DiMarino D, Buzzi A, et al. CYFIP1 coordinates mRNA translation and cytoskeleton remodeling to ensure proper dendritic Spine formation. Neuron. 2013;79(6). doi:10.1016/j.neuron.2013.06.039

20. Thomas P, Smart TG. HEK293 cell line: A vehicle for the expression of recombinant proteins. J Pharmacol Toxicol Methods. 2005;51(3 SPEC. ISS.). doi:10.1016/j.vascn.2004.08.014

21. Vidal RL, Valenzuela JI, Luján R, Couve A. Cellular and subcellular localization of Marlin-1 in the brain. BMC Neurosci. 2009;10. doi:10.1186/1471-2202-10-37

22. Luther DC, Jeon T, Goswami R, Nagaraj H, Kim D, Lee YW, et al. Protein Delivery: If Your GFP (or Other Small Protein) Is in the Cytosol, It Will Also Be in the Nucleus. Bioconjugate Chemistry. 2021. doi:10.1021/acs.bioconjchem.1c00103

23. Bräuer M, Zich MT, Önder K, Müller N. The influence of commonly used tags on structural propensities and internal dynamics of peptides. Monatshefte für Chemie - Chemical Monthly. 2019 May 29;150(5):913–25. doi:10.1007/s00706-019-02401-x

24. Pathania M, Davenport EC, Muir J, Sheehan DF, López-Doménech G, Kittler JT. The autism and schizophrenia associated gene CYFIP1 is critical for the maintenance of dendritic complexity and the stabilization of mature spines. Transl Psychiatry. 2014;4. doi:10.1038/tp.2014.16

25. Davenport EC, Szulc BR, Drew J, Taylor J, Morgan T, Higgs NF, et al. Autism and Schizophrenia-Associated CYFIP1 Regulates the Balance of Synaptic Excitation and Inhibition. Cell Rep. 2019;26(8). doi:10.1016/j.celrep.2019.01.092

26. Martin JG, Martin ER, Takamura N, Harlow CE, Bamford RA, Smith RG, et al. Exploring the functions of JAKMIP1 in neuronal IL-6/STAT3 signaling and its relevance to chromosome 15q-duplication syndrome [Internet]. 2025. Available from: http://biorxiv.org/lookup/doi/10.1101/2025.10.21.683757 doi:10.1101/2025.10.21.683757

27. Willems J, de Jong APH, Scheefhals N, Mertens E, Catsburg LAE, Poorthuis RB, et al. Orange: A CRISPR/Cas9-based genome editing toolbox for epitope tagging of endogenous proteins in neurons. PLoS Biol. 2020;18(4). doi:10.1371/journal.pbio.3000665

